# Colocalization of IQGAP1 with DLC-1 and PLC-δ1; Its potential role in coronary spasm

**DOI:** 10.1101/526152

**Authors:** Makoto Tanaka, Tomohiro Osanai, Yoshimi Homma, Kenji Hanada, Ken Okumura, Hirofumi Tomita

**Author notes:** Correspondence to: Makoto Tanaka, PhD, Department of Stroke and Cerebrovascular Medicine, Hirosaki University Graduate School of Medicine, 5 Zaifu-cho, Hirosaki, 036-8562 Japan.

## Abstract

Phospholipase C (PLC)-δ1, activated by p122RhoGTPase-activating protein (GAP)/deleted in liver cancer-1 (p122RhoGAP/DLC-1), contributes to the coronary spastic angina (CSA) pathogenesis. The present study aims to further investigate the p122RhoGAP/DLC-1 protein. We examined molecules assisting this protein and identified a scaffold protein—IQ motif-containing GTPase-activating protein 1 (IQGAP1). IQGAP1-C binds to the steroidogenic acute regulatory-related lipid transfer (START) domain of p122RhoGAP/DLC-1, and PLC-δ1 binds to IQGAP1-N, forming a complex. In fluorescence microscopy, small dots of PLC-δ1 created fine linear arrays like microtubules, and IQGAP1 and p122RhoGAP/DLC-1 were colocated in the cytoplasm with PLC-δ1. Ionomycin induced the raft recruitment of the PLC-δ1, IQGAP1, and p122RhoGAP/DLC-1 complex by translocation to the plasma membrane (PM), indicating the movement of this complex is along microtubules with the motor protein kinesin. Moreover, the IQGAP1 protein was elevated in skin fibroblasts obtained from patients with CSA, and it enhanced the PLC activity and peak intracellular calcium concentration in response to acetylcholine. IQGAP1, a novel stimulating protein, forms a complex with p122RhoGAP/DLC-1 and PLC-δ1 that moves along microtubules and enhances the PLC activity.

## Introduction

Coronary artery spasm, an abnormal contraction of the epicardial coronary artery responsible for myocardial ischemia, is essential in the pathogenesis of Prinzmetal variant angina (Oliva et al., 1973, Yasue et al., 1974), myocardial infarction with nonobstructive coronary arteries (Pasupathy et al., 2015), malignant ventricular arrhythmias (Kaku et al., 2015, Sanna et al., 2009), and the other acute coronary syndromes (Okumura et al., 1996, Oliva et al., 1977)—all of which can lead to sudden death. In certain Japanese studies on coronary spastic angina (CSA), basal vasomotor tone and constrictive response to various stimuli within the coronary arteries were enhanced (Hoshio et al., 1989, Kaski et al., 1986, Kuga et al., 1993, Okumura et al., 1996). These findings attributed the hyperactivity of the coronary artery smooth muscle to intracellular and/or postreceptorial mechanisms (Lanza et al., 2011).

Phospholipase C (PLC) correlates with the contraction of coronary arteries and is a vital molecule in the intracellular calcium regulation. PLC hydrolyzes phosphatidylinositol 4,5-bisphosphate (PIP_2_) to produce inositol 1,4,5-trisphosphate (IP3) and diacylglycerol. IP3 mobilizes Ca^2+^ from the intracellular stores and exhibits rapid contraction of the vascular smooth muscle (Korn et al., 1988), whereas diacylglycerol activates protein kinase C and triggers the sustained muscle contraction via a Ca^2+^-independent mechanism (Ito et al., 1994). Previously, we have reported the enhanced PLC activity in cultured skin fibroblasts obtained from patients with CSA and determined that a major PLC isozyme in the membrane fraction was the δ1 isoform, which is more sensitive to Ca^2+^ than other isozymes (Okumura et al., 2000). Furthermore, we have demonstrated the presence of a G to A mutation at nucleotide position 864 in PLC-δ1 in patients with CSA, accompanied by the amino acid (aa) replacement of arginine 257 to histidine (R257H), which markedly enhanced the PLC enzymatic activity in the physiological range of the intracellular free calcium concentration ([Ca^2+^]_i_) (Nakano et al., 2002). To elucidate its role in coronary spasm, we created mice that overexpressed the variant PLC-δ1 (R257H) under the control of the mouse α-SMA promoter and illustrated that the elevated PLC-δ1 activity enhanced coronary vasomotility, as observed in patients with CSA (Shibutani et al., 2012).

A study has demonstrated that p122RhoGTPase-activating protein (GAP)/deleted in liver cancer-1 (p122RhoGAP/DLC-1), cloned as a PLC-δ1–interacting protein from a rat brain expression library, exhibits a specific GTPase-activating protein (GAP) activity on Rho, augmenting the PIP_2_ hydrolyzing activity of PLC-δ1 *in vitro* (Homma et al., 1995). Further, this rat p122RhoGAP/DLC-1 was reportedly the ortholog of human p122RhoGAP/DLC-1, with an aa sequence identity of 93% (Durkin et al., 2007, Yuan et al., 1998, Yuan et al., 2003). In addition, p122RhoGAP/DLC-1 is acknowledged as a tumor suppressor and is downregulated in several malignant cancer types, including colorectal, breast, prostate, and liver cancer (Liao et al., 2008). Previously, we have reported that the p122RhoGAP/DLC-1 protein expression in cultured skin fibroblasts obtained from patients with CSA was upregulated three-fold compared with that in control fibroblasts, and the p122RhoGAP/DLC-1 overexpression increased [Ca^2+^]_i_ in response to acetylcholine (ACh) (Murakami et al., 2010). Furthermore, we created mice that overexpressed p122RhoGAP/DLC-1 and illustrated that the coronary spasm was induced by injecting ergometrine into the jugular vein (Kinjo et al., 2015). These findings revealed that the p122RhoGAP/DLC-1 upregulation in coronary arteries is attributable to the coronary spasm pathogenesis associated with human CSA. Nevertheless, the regulatory mechanisms and physiological functions of p122RhoGAP/DLC-1 and PLC-δ1 remain partially understood. The present study aims to identify a novel protein that interacts with p122RhoGAP/DLC-1 and PLC-δ1 and elucidate its role in the CSA pathogenesis.

## Results

### ▪ p122RhoGAP/DLC-1 protein expression in cultured fibroblasts

We detected the p122RhoGAP/DLC-1 protein using 4%–20% gradient sodium dodecyl sulfate-polyacrylamide gel electrophoresis (SDS–PAGE) gel in a single band around 122 kDa, and its expression was increased in patients with CSA as anticipated (Fig 1A). Remarkably, we detected an unknown band around 200 kDa (150–250 kDa) above the p122RhoGAP/DLC-1 protein band in patients with CSA after prolonged exposure; as this unknown band was detected with a specific antibody against p122RhoGAP/DLC-1, we assumed it to be a heterodimer of p122RhoGAP/DLC-1 and its binding protein. In addition, the molecular weight of p122RhoGAP/DLC-1–binding protein was estimated to be approximately 80 kDa (difference between 200 and 122 kDa). Furthermore, the expression of this unknown band was higher in patients with CSA than that in control subjects.

**Fig 1.**
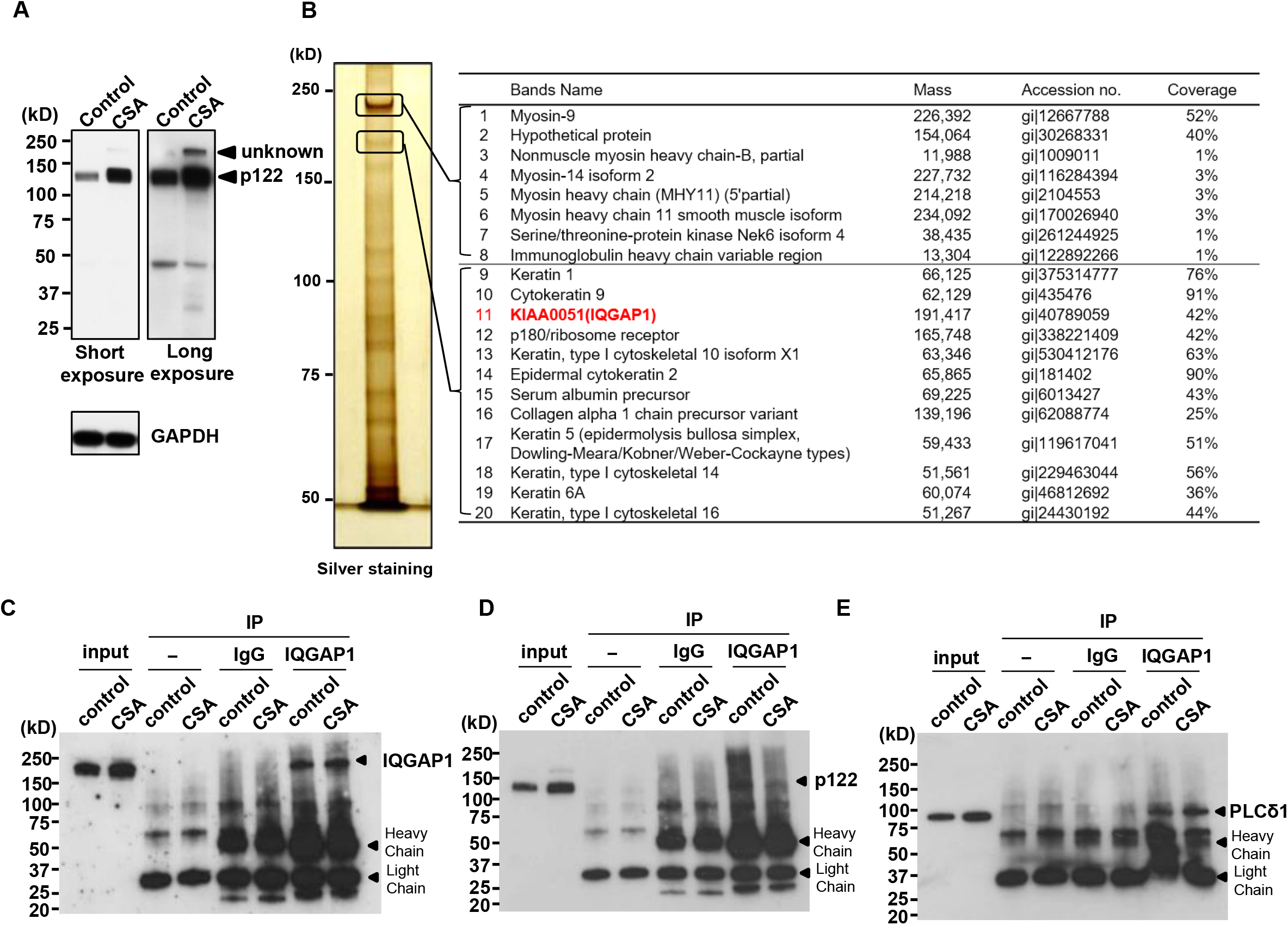
The identification of p122RhoGAP/DLC-1–binding protein and interaction of IQGAP1 with p122RhoGAP/DLC-1 and PLC-δ1 in skin fibroblasts. A, The human p122RhoGAP/DLC-1 and GAPDH protein expression using 4%–20% gradient sodium dodecyl sulfate–polyacrylamide gel electrophoresis (SDS–PAGE) gel in skin fibroblasts of control subjects (*n* = 3) and patients with coronary spastic angina (CSA; *n* = 6). To detect unknown signal intensities, Western blotting was additionally exposed for a prolonged duration (long exposure, 10 min). B, Skin fibroblasts cell lysates obtained from CSA were immunoprecipitated using an anti-p122RhoGAP/DLC-1 mouse monoclonal antibody, fractionated by 7.5% SDS– PAGE, and detected with silver staining followed by time of flight mass spectrometry (TOF-MS) analysis of p122RhoGAP/DLC-1 peptide-interacting proteins. C, D, and E, Cell lysates immunoprecipitated with an anti-IQGAP1 mouse monoclonal antibody or with and without normal mouse IgG. C, The IQGAP1 IP elute is resolved on SDS–PAGE and immunoblotted using an IQGAP1 antibody. D, The IQGAP1 IP elute is resolved on SDS–PAGE and immunoblotted using a p122RhoGAP/DLC-1 antibody. E, The IQGAP1 IP elute is resolved on SDS–PAGE and immunoblotted using a PLC-δ1 antibody. IP, immunoprecipitation.

### ▪ Identification of p122RhoGAP/DLC-1–binding proteins

We performed immunoprecipitation assay of p122Rho–GAP/DLC-1 using CSA skin fibroblasts to analyze the unknown protein mentioned in the section above. Electrophoresis was carried out using 7.5% gel to expand the range of 150–250 kDa. Several proteins bound specifically to p122RhoGAP/DLC-1 were detected by an immunoprecipitation assay with skin fibroblast lysate and visualized by silver staining (Fig 1B). Of them, proteins around 200 kDa (150–250 kDa) were characterized by using the time of flight mass spectrometry (TOF-MS) method using TripleTOF 5600 (AB Sciex Pte. Ltd.) and then by entering the resulting data into a search of the NCBI database by using ProteinPilot™ Software 4.5 (AB Sciex Pte. Ltd.). Surprisingly, p122RhoGAP/DLC-1 was not detected in the band around 200 kDa being relevant to the heterodimer (Fig 1B, proteins 1–20), indicating that the unknown band is not a heterodimer of p122RhoGAP/DLC-1, but its binding protein. Proteins 1–8 identified in the upper band were myosins, the principal protein that constitutes myofibrils, and only two proteins exhibited high coverage. In proteins 9–20 identified in the lower band, the main identified protein was keratin (around 51–66 kDa). Of note, protein of about 80 kDa was not detected in upper and lower bands. Hence, we confirmed the unknown band to be neither a heterodimer of p122RhoGAP/DLC-1 nor its binding protein. Rather, KIAA0051 [IQ motif-containing GAP 1 (IQGAP1)] of 191 kDa in protein 11, which shared similar characteristics to p122RhoGAP/DLC-1, was detected as an unknown band protein and as a candidate for the p122RhoGAP/DLC-1–binding protein (Fig 1B, right), indicating that the unknown band highly expressed in CSA fibroblasts is not specific for the anti-p122RhoGAP/DLC-1 antibody but p122RhoGAP/DLC-1–binding protein. Furthermore, p122RhoGAP/DLC-1 is an intracellular protein that forms part of the cytoskeleton, such as myosin and keratin.

### ▪ Interaction of IQGAP1 with p122RhoGAP/DLC-1 and PLC-δ1 in skin fibroblasts

The mass spectrometry analysis revealed that IQGAP1 interacts with p122RhoGAP/DLC-1; using immunoprecipitation (Woeller et al., 2015), we identified this interaction in skin fibroblasts obtained from control subjects and patients with CSA. The IQGAP1-specific immunoprecipitation elute was immunoblotted with p122RhoGAP/DLC-1 and PLC-δ1 antibodies (Fig 1D and E). We detected the bands corresponding to these molecules in an IQGAP1 immunoprecipitate, establishing the correlation of p122RhoGAP-DLC-1 and PLC-δ1 with IQGAP1. Of note, PLC-δ1-like bands on the lanes of no anti-IQGAP1 antibody or IgG in Fig 1E were nonspecific (molecular weight of PLC-δ1-like bands is higher than that of PLC-δ1).

### ▪ Subcellular localization of p122RhoGAP/DLC-1, IQGAP1, and PLC-δ1 in skin fibroblasts

As the immunoprecipitation study illustrated the interaction of IQGAP1 with p122RhoGAP/DLC-1 and PLC-δ1, we hypothesized that these proteins would be colocalized in skin fibroblasts. As shown in Fig 2A (a, c, and e), IQGAP1 and p122RhoGAP/DLC-1 were primarily localized in the nucleus and, to a small extent, in the cytoplasm. To elucidate these points, we chose an extended flat area of the cell and magnified images of the two proteins. IQGAP1 and p122RhoGAP/DLC-1 were arranged in small dots and created a structure of linear arrays in the cytoplasm (Fig 2A, b and d). Upon merging these images, the linear arrays of IQGAP1 and p122RhoGAP/DLC-1 were colocalized in the cytoplasm (Fig 2A, f).

**Fig 2.**
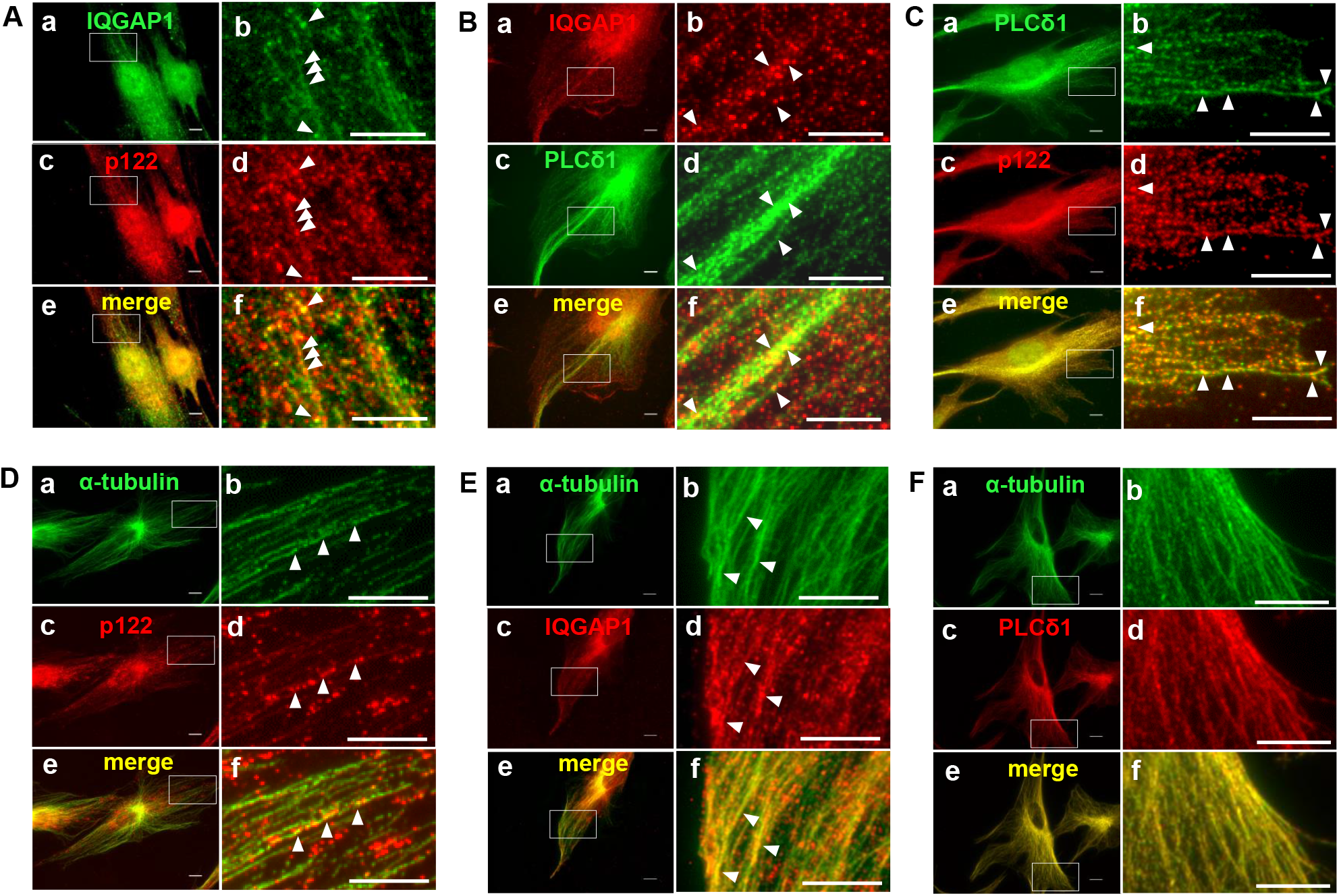
The subcellular localization of IQGAP1, p122RhoGAP/DLC-1, and PLC-δ1 and location of p122RhoGAP/DLC-1, IQGAP1, and PLC-δ1 along microtubules in coronary spastic angina (CSA) skin fibroblasts. A, Double immunostaining of IQGAP1 and p122 in skin fibroblasts. The localization of IQGAP1 is indicated by the green fluorescence of Alexa Fluor 488 (a and b). The localization of p122 is indicated by the red fluorescence of Texas Red-X (c and d). Merged images are shown (e and f). One flat, extended area is magnified (b, d, and f). B, Double immunostaining of IQGAP1 and PLC-δ1 in skin fibroblasts. The localization of IQGAP1 is indicated by the red fluorescence of Alexa Fluor 594 (a and b). The localization of PLC-δ1 is indicated by the green fluorescence of FITC (c and d). Merged images are shown (e and f). One flat, extended area is magnified (b, d, and f). C, Double immunostaining of PLC-δ1 and p122 in skin fibroblasts. The localization of PLC-δ1 is indicated by the green fluorescence of FITC (a and b). The localization of p122 is indicated by the red fluorescence of Texas Red-X (c and d). Merged images are shown (e and f). One flat, extended area is magnified (b, d, and f). D, Double immunostaining of α-tubulin and p122 in skin fibroblasts. The localization of α-tubulin is indicated by the green fluorescence of Alexa Fluor 488 (a and b). The localization of p122 is indicated by the red fluorescence of Texas Red-X (c and d). Merged images are shown (e and f). One flat, extended area is magnified (b, d, and f). E, Double immunostaining of α-tubulin and IQGAP1 in skin fibroblasts. The localization of α-tubulin is indicated by the green fluorescence of Alexa Fluor 488 (a and b). The localization of IQGAP1 is indicated by the red fluorescence of Alexa Fluor 594 (c and d). Merged images are shown (e and f). One flat, extended area is magnified (b, d, and f). F, Double immunostaining of α-tubulin and PLC-δ1 in skin fibroblasts. The localization of α-tubulin is indicated by the green fluorescence of Alexa Fluor 488 (a and b). The localization of PLC-δ1 is indicated by the red fluorescence of Texas Red-X (c and d). Merged images are shown (e and f). One flat, extended area is magnified (b, d, and f). Scale bars, 10 μm.

IQGAP1 was primarily localized in the nucleus as small dots and created slight linear arrays in the cytoplasm (Fig 2B, a). PLC-δ1 was primarily detected in the cytoplasm as small dots and only to a small extent in the nucleus (Fig 2B, c). The small dots of PLC-δ1 created fine linear arrays, similar to microtubules, which were colocalized in the cytoplasm with the linear arrays of IQGAP1 (Fig 2B, e). Once more, we chose an extended flat area of the cell and magnified the images; merging these revealed that the small dots of IQGAP1 were located along the arrays of PLC-δ1 (Fig 2B, b, d, and f).

Small dots of PLC-δ1 and p122RhoGAP/DLC-1 created fine linear arrays like microtubules (Fig 2C, a–d); these were colocalized in the cytoplasm. Merging magnified images of an extended flat area of the cell revealed that the linear arrays of dots of p122RhoGAP/DLC-1 were located along the arrays of PLC-δ1 (Fig 2C, f).

### ▪ Location of p122RhoGAP/DLC-1, IQGAP1, and PLC-δ1 along microtubules

As explained above, p122RhoGAP/DLC-1, IQGAP1, and PLC-δ1 displayed linear arrays and, perhaps, could be involved in the transport system along microtubules; this observation led us to compare the distributions of p122RhoGAP/DLC-1, IQGAP1, PLC-δ1, and the microtubule network. Accordingly, we double-immunostained p122RhoGAP/DLC-1, IQGAP1, or PLC-δ1 in skin fibroblasts, as well as α-tubulin for the microtubule network, and ascertained their distributions by fluorescence microscopy. The α-tubulin created a fine network of microtubules (Fig 2D, a, E, a, and F, a). We observed p122RhoGAP/DLC-1 in the form of cytoplasmic bodies (Fig 2D, c). For elucidation, we chose an extended flat area of the cell and magnified images of the two proteins. Small dots of α-tubulin and p122RhoGAP/DLC-1 formed fine linear arrays (Fig 2D, b and d). Merging the images displayed an abundance of p122RhoGAP/DLC-1 cytoplasmic bodies along the microtubule network (Fig 2D, f). Notably, we observed p122RhoGAP/DLC-1 cytoplasmic bodies to be partially located along microtubules; similarly, IQGAP1 and PLC-δ1 were observed as cytoplasmic bodies (Fig 2E, c and F, c). For clarification, we selected an extended flat area of the cell and magnified images of the two proteins. Small dots of IQGAP1 formed fine linear arrays (Fig 2E, d), which in merged images were found to be located along microtubules (Fig 2E, f). Furthermore, small dots of PLC-δ1 formed a fine network like microtubules (Fig 2F, f). Remarkably, α-tubulin and PLC-δ1 were almost colocalized in the cytoplasm. These findings established that p122RhoGAP/DLC-1, IQGAP1, and PLC-δ1 were located along microtubules.

### ▪ IQGAP1 protein expression in cultured fibroblasts

After 16-h starvation, we scraped cultured fibroblasts to assess the IQGAP1 protein expression. The anti-IQGAP1 antibody was specific to IQGAP1 (Fig 3A), and IQGAP1 was strongly detected as a single band of 195 kDa just above the 150-kDa marker. Compared with control subjects, the IQGAP1-to-GAPDH protein ratio was significantly higher by 1.4 ± 0.2 times in patients with CSA (*P* < 0.01; Fig 3B).

**Fig 3.**
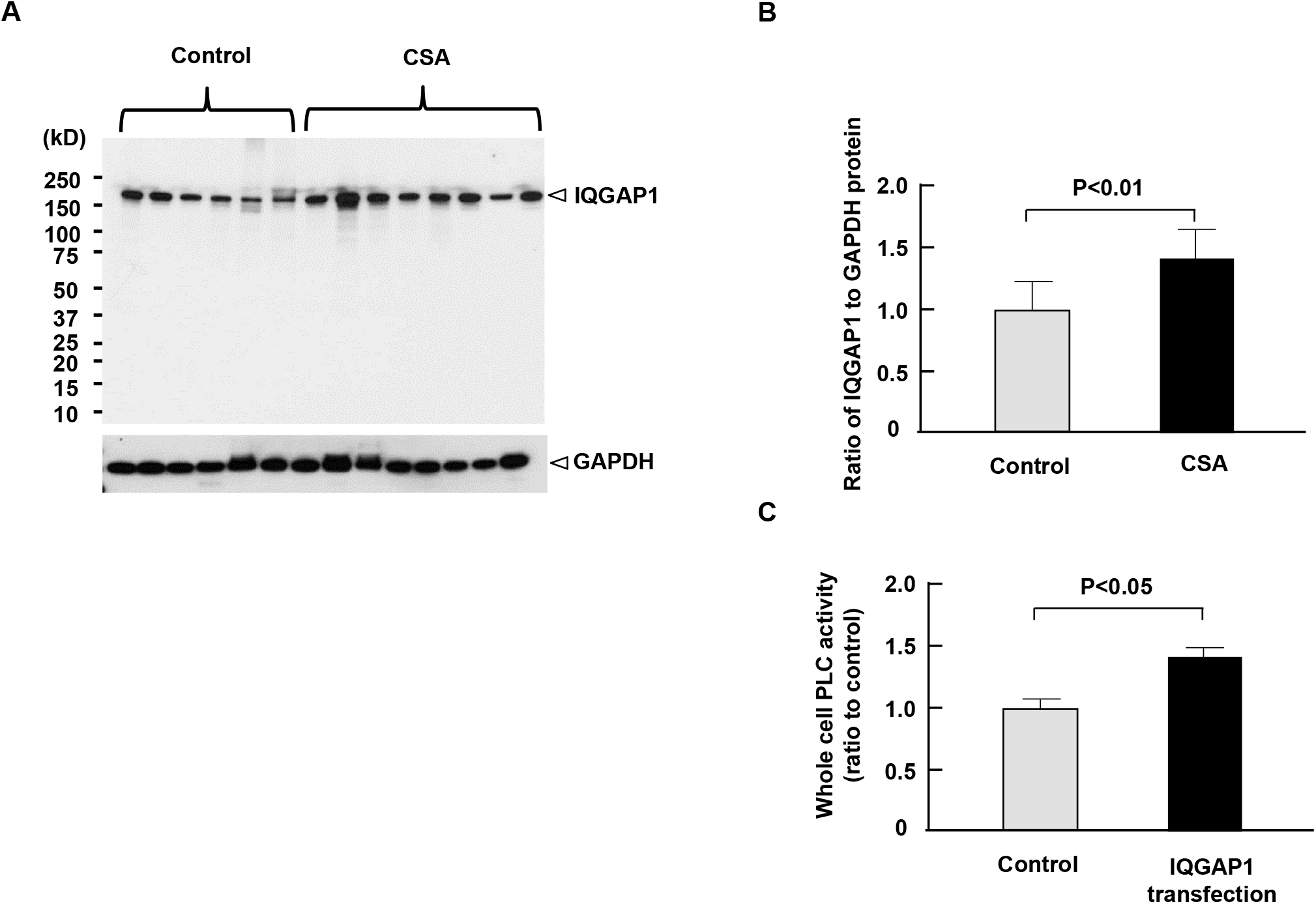
The IQGAP1 protein expression in cultured fibroblasts. A, The human p122RhoGAP/DLC-1 and GAPDH protein expression in skin fibroblasts of control subjects (*n* = 6) and patients with coronary spastic angina (CSA; *n* = 8). B, The IQGAP1 protein expression in skin fibroblasts normalized by human GAPDH in patients with CSA and control subjects. C, The phospholipase C (PLC) activity in HEK293 cells with and without IQGAP1 transfection (*n* = 3).

### ▪ PLC activity

In this study, the PLC enzymatic activity was increased by 1.4 ± 0.05 times in IQGAP1-transfected human embryonic kidney (HEK) 293 cells compared with those transfected with an empty vector (*n* = 3, *P* < 0.05; Fig 3C).

### ▪ Impact of IQGAP1 transfection on the [Ca^2+^]_i_ response to ACh

We assessed the impact of the IQGAP1 overexpression on the ACh-induced increase in [Ca^2+^]_i_ using HEK293 cells. Fig 4A shows representative waveforms of [Ca^2^]_i_ in HEK293 cells after the ACh administration at 10^−4^ M. The ACh-induced increment in [Ca^2+^]_i_ was augmented by the IQGAP1 overexpression, which was established by Western blotting as a single immunoreactive band at 195 kDa (Fig 4D). The IQGAP1-to-GAPDH ratio in IQGAP1-overexpressed cells was 1.95 ± 0.15 times (*n* = 5, *P* < 0.01) compared with that in cells transfected with an empty vector rather than IQGAP1 (Fig 4D). The [Ca^2+^]_i_ levels at the baseline were 14 ± 9 nM without the IQGAP1 overexpression and 53 ± 34 nM with the IQGAP1 overexpression (Fig 4B, *n* = 5, *P* < 0.05). In addition, the ACh-induced peak increase in [Ca^2+^]_i_ was higher in cells with the IQGAP1 overexpression than those without the IQGAP1 overexpression (134 ± 23 nM vs. 95 ± 10 nM, *n* = 5, *P* < 0.01; Fig 4C).

**Fig 4.**
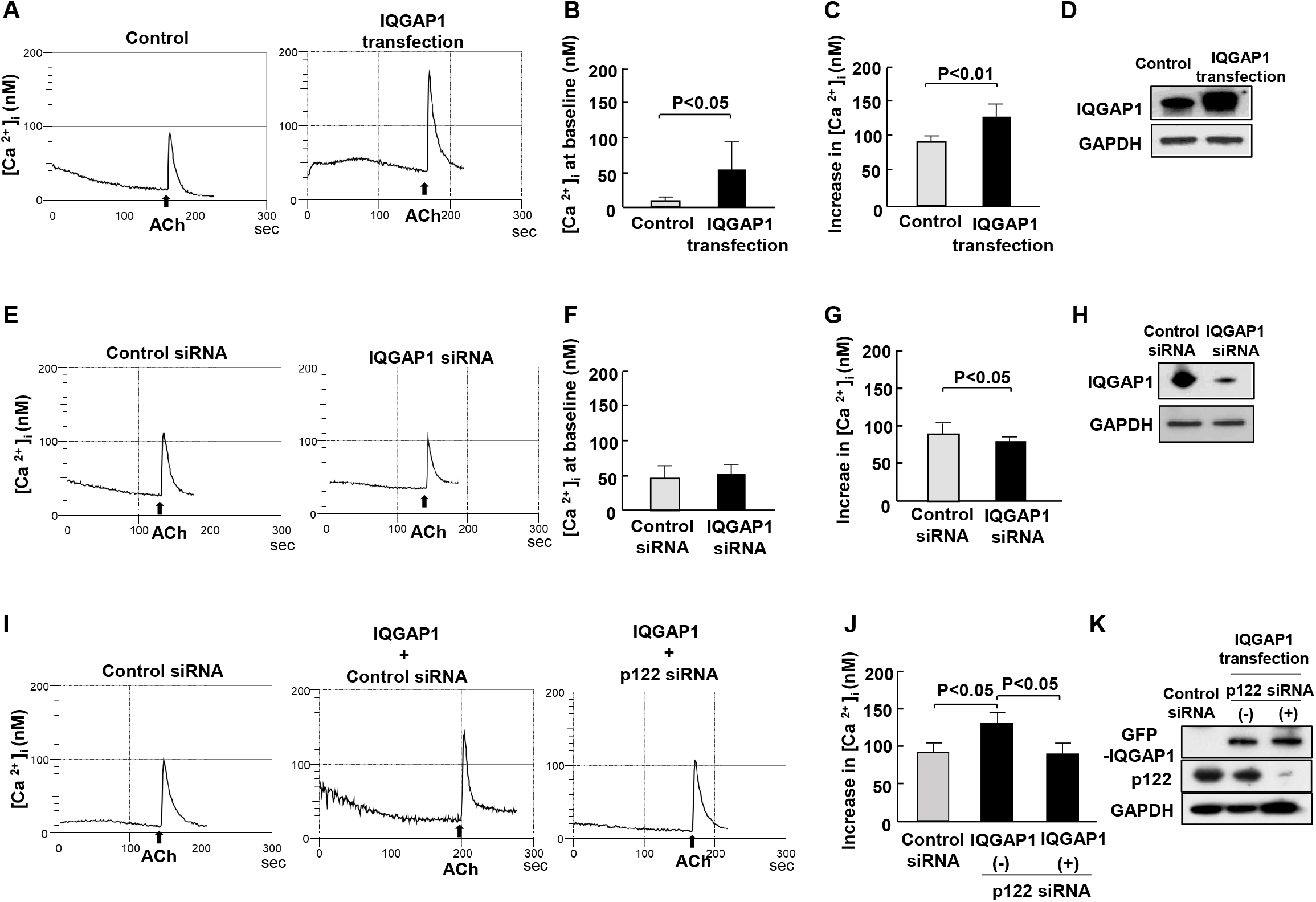
The effect of IQGAP1 transfection on the response of [Ca^2+^]i to acetylcholine (ACh), and phospholipase C (PLC) activity. A, E, and I, The representative wave forms of [Ca^2+^]i. B, The intracellular calcium concentration [Ca^2+^]i level at the baseline after 10
^−4^ M ACh in HEK293 cells with and without IQGAP1 (*n* = 5). C, The peak increase in [Ca^2+^]_i_ from the baseline after 10^−4^ M ACh in HEK293 cells with and without IQGAP1 (*n* = 5). D, The IQGAP1 and GAPDH protein expression in HEK293 cells with and without IQGAP1 transfection. F, The intracellular calcium concentration [Ca^2+^]i level at the baseline after 10^−4^ M ACh in HEK293 cells with and without IQGAP1 siRNA (*n* = 5). G, The peak increase in [Ca^2+^]i from the baseline after 10^−4^ M ACh in cells with and without IQGAP1 siRNA (*n* = 5). H, The IQGAP1 and GAPDH protein expression in HEK293 cells with and without IQGAP1 siRNA transfection. J, The peak increase in [Ca^2+^]i from the baseline after 10^−4^ M ACh in cells with and without IQGAP1 or p122 siRNA transfection (*n* = 5). K, The GFP-IQGAP1, p122RhoGAP/DLC-1, and GAPDH protein expression in HEK293 cells with and without IQGAP1 or p122 siRNA transfection.

### ▪ Effect of IQGAP1 siRNA transfection on the response of [Ca^2+^]i to ACh

Next, we investigated the impact of IQGAP1 siRNA on the ACh-induced increment in [Ca^2+^]_i_ in HEK293 cells. The IQGAP1-to-GAPDH ratio was 79.7% ± 2.5% (*n* = 5, *P* < 0.01) lower in IQGAP1 siRNA–transfected cells than those transfected with the control siRNA vector (Fig 4H). The [Ca^2+^]_i_ levels at the baseline were similar between IQGAP1 siRNA-transfected cells and the control siRNA vector (Fig 4F, *n* = 5). The peak increase in [Ca^2+^]_i_ was marginally lower in IQGAP1 siRNA-transfected cells than in those transfected with the control siRNA vector (77 ± 4 nM vs. 95 ± 13 nM, *n* = 5, *P* < 0.05; Fig 4E and G).

### ▪ Impact of the IQGAP1 overexpression in HEK293 cells with and without the p122RhoGAP/DLC-1 knockdown

In this study, HEK293 cells were cotransfected with IQGAP1 and p122RhoGAP/DLC-1 siRNA to disrupt the complex, and we measured the increase in [Ca^2+^]_i_ with ACh at 10 ^−4^ M. The p122RhoGAP/DLC-1-to-GAPDH ratio was 95.0% ± 2.7% (*n* = 5, P < 0.01) lower in cells cotransfected with IQGAP1 and p122RhoGAP/DLC-1 siRNA than in those transfected with the control siRNA vector (Fig 4K). The IQGAP1 overexpression enhanced the ACh-induced increase in [Ca^2+^]_i_ (Fig 4I and J). Nevertheless, in HEK293 cells cotransfected with IQGAP1 and p122RhoGAP/DLC-1 siRNA, the ACh-induced increase in [Ca^2+^]_i_ was suppressed from 135 ± 18 nM to 94 ± 15 nM, being relevant to a level in control cells transfected with the control siRNA vector (95 ± 13 nM; Fig 4I and J).

### ▪ *In vitro* interaction between IQGAP1 and p122RhoGAP/DLC-1

We conducted an *in vitro* interaction assay and attempted to narrow down the p122RhoGAP/DLC-1 region essential for IQGAP1 binding to confirm the interaction between IQGAP1 and p122RhoGAP/DLC-1 observed in immunoprecipitation. In addition, we conducted pull-down experiments with the p122RhoGAP/DLC-1 deletion mutants (aa 1–1091, 1–546, 1–801, and 547–1091; Fig 5B) expressed in bacteria with glutathione S-transferase (GST)-fused IQGAP1-N and IQGAP1-C (Fig 5A). The N- or C-terminal halves of IQGAP1 (IQGAP1-N, aa 1–863, and IQGAP1-C, aa 746–1657) were expressed as GST fusion proteins and purified from the bacteria (Fig 5A). Silver staining revealed that the sizes of GST-IQGAP1-N and GST-IQGAP1-C fusion proteins were 124 and 131 kDa, respectively, as anticipated (Fig 5D). We observed that p122RhoGAP/DLC-1 (1–1091) and p122RhoGAP/DLC-1 (547–1091) could be precipitated with GST-IQGAP1-C but not with GST-IQGAP1-N (Fig 5E). Furthermore, p122RhoGAP/DLC-1 (1–546) and p122RhoGAP/DLC-1 (1–801) could not be precipitated with GST-IQGAP1-C and -N (Fig 5E). These findings suggested that IQGAP1-C interacts through the region covering residues 802-1091 of the START domain of p122RhoGAP/DLC-1.

**Fig 5.**
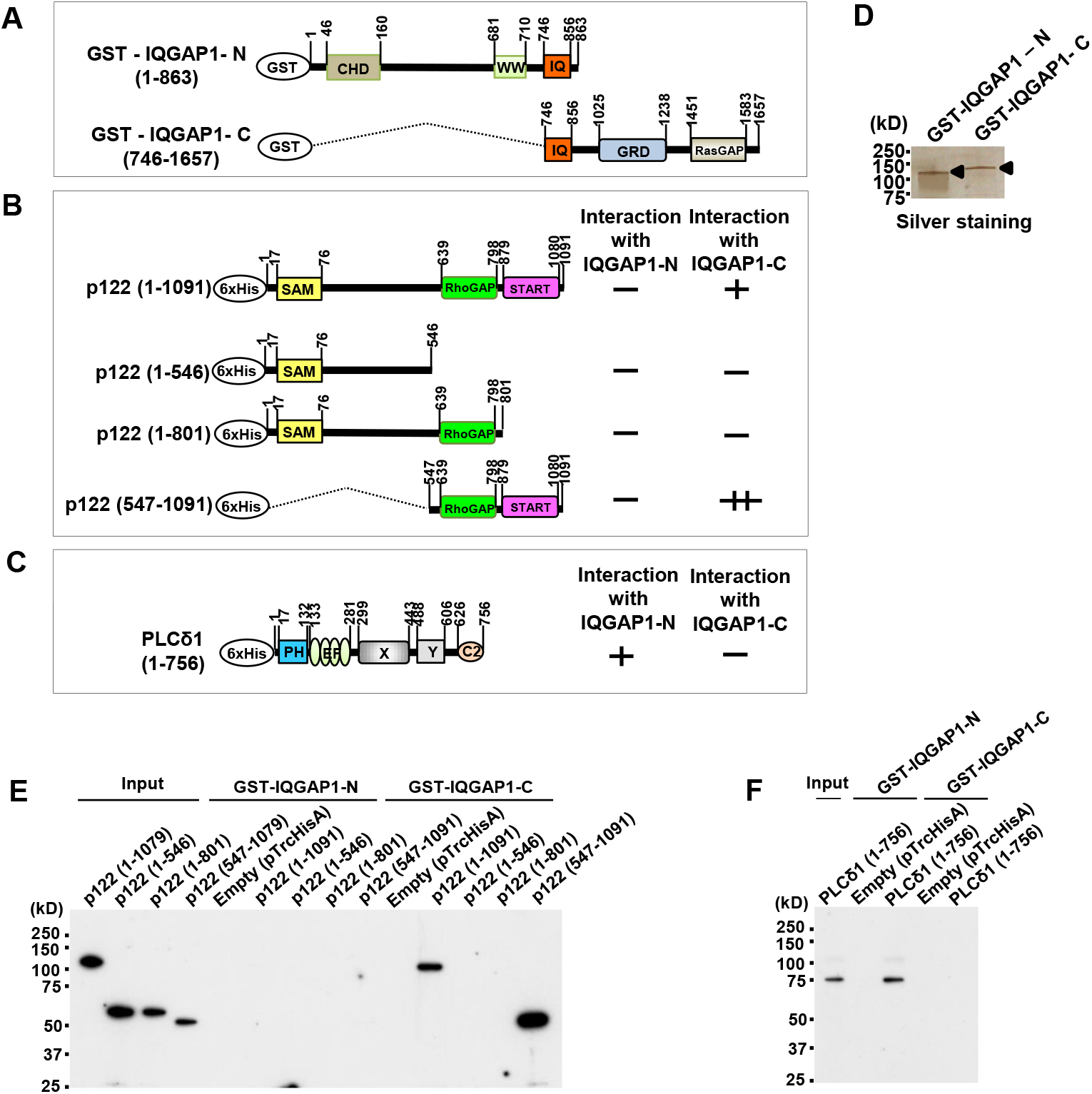
The *in vitro* interaction between IQGAP1, p122RhoGAP/DLC-1, and PLC-δ1. A, The schematic presentation of the domain organization of the IQGAP1. The numbers above each construct refer to the amino acid (aa) numbers. CHD, calponin homology domain; WW, tryptophan-containing proline-rich motif-binding region; IQ repeats, IQGAP-specific repeats; GRD, RasGAP-related domain; RasGAP, RasGAP C-terminal domain. B, The schematic presentation of the domain organization of the p122RhoGAP/DLC-1. The numbers above each construct refer to the aa numbers. SAM, sterile α motif; RhoGAP, RhoGAP domain; START, steroidogenic acute regulatory-related lipid transfer. The interaction between IQGAP1-C (746–1657) and truncated p122RhoGAP/DLC-1 is summarized. C, The schematic presentation of the domain organization of the PLC-δ1. The numbers above each construct refer to the aa numbers. PH, pleckstrin homology domain; EF, EF-hand motif tandem potential Ca^2+^ binding motifs; X and Y, central catalytic domains; C2, Ca^2+^-dependent phospholipid binding domain. The interaction between IQGAP1-N (1–863) and truncated PLC-δ1 is summarized. D, The analysis of purified GST-IQGAP1-N and GST-IQGAP1-C fusion proteins on sodium dodecyl sulfate–polyacrylamide gel electrophoresis (SDS–PAGE) stained with sliver staining. E, Pull-down experiments with various deletion mutants of p122 (1–1079), p122 (1– 546), p122 (1–801), and p122 (547–1079) expressed in bacteria with glutathione S-transferase (GST)-fused IQGAP1-N and IQGAP1-C are performed. F, Pull-down experiments with various deletion mutants of PLC-δ1 (1–756) expressed in bacteria with GST-fused IQGAP1-N and IQGAP1-C are performed.

### ▪ *In vitro* interaction between IQGAP1 and PLC-δ1

Likewise, we conducted an *in vitro* interaction assay, conducting pull-down experiments with PLC-δ1 (1–756) (Fig 5C) expressed in bacteria with GST-fused IQGAP1-N and IQGAP1-C to confirm the interaction between IQGAP1 and PLC-δ1 observed in skin fibroblasts (Fig 5A). PLC-δ1 (1–756) could be precipitated with GST-IQGAP1-N but not with GST-IQGAP1-C (Fig 5F), indicating that PLC-δ1 interacts through the region covering residues 1–745 of the CHD and WW domain of IQGAP1. However, further studies are warranted to elucidate the exact binding domain of PLC-δ1 to IQGAP1.

### ▪ Recruitment of PLC-δ1, IQGAP1, and p122RhoGAP/DLC-1 to the PM

p122RhoGAP/DLC-1 is a PLC-δ1–interacting protein localized in lipid rafts in fibroblastic and epithelial cell lines. Reportedly, the interaction between PLC-δ1 and p122RhoGAP/DLC-1 is enhanced by treating PC12 cells with carbamylcholine (Yamaga et al., 2008). By investigating the GFP–PLC-δ1-PH expression and analyzing its involvement in the endogenous IQGAP1 localization by immunostaining, Choi et al. (2013) revealed that in the optimal amount of PLC-δ1-PH domain, endogenous IQGAP1 partially colocalized with GFP–PLC-δ1-PH in MDA-MB-231 cell lines.

In this study, we investigated whether IQGAP1, p122RhoGAP/DLC-1, and PLC-δ1 were recruited to the PM after stimulating skin fibroblasts with ionomycin, a Ca^2+^ ionophore. We treated skin fibroblasts with or without ionomycin at 10^−5^ M for 5 min and analyzed IQGAP1, p122RhoGAP/DLC-1, and PLC-δ1 using Western blotting in the total cell lysates (TCL), cytosol (CY), and the PM fractions. No change was noted during this process in the amounts of IQGAP1, p122RhoGAP/DLC-1, and PLC-δ1 in the TCL (Fig 6A), nor was there a change in the amount of IQGAP1 in the CY (Fig 6A). However, we observed an apparent increase of IQGAP1 in the PM after the ionomycin treatment (Fig 6A). Similarly, the amounts of p122RhoGAP/DLC-1 and PLC-δ1 in the CY decreased after the ionomycin treatment but increased in the PM. These findings suggested that IQGAP1, p122RhoGAP/DLC-1, and PLC-δ1 in skin fibroblasts were recruited from the CY to the PM.

**Fig 6.**
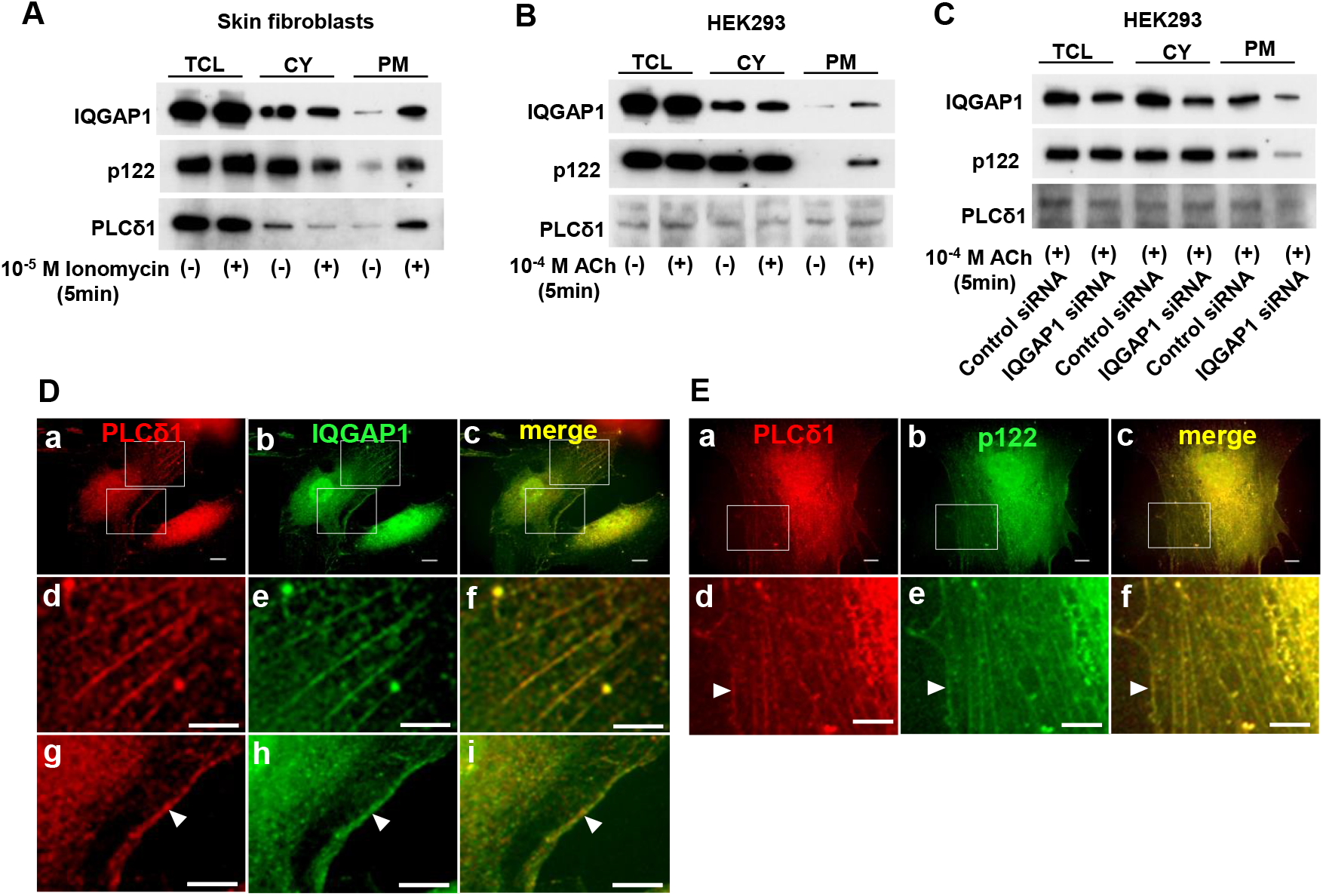
The recruitment of IQGAP1, p122RhoGAP/DLC-1, and PLC-δ1 to the plasma membrane. A, Western blotting displaying the IQGAP1, p122RhoGAP/DLC-1, and PLC-δ1 protein level in the total cell lysates (TCL), cytosol (CY), and plasma membrane (PM) of skin fibroblasts cells treated with or without ionomycin 10^−5^ M for 5 min. B, Western blotting displaying the IQGAP1, p122RhoGAP/DLC-1, and PLC-δ1 protein level in the TCL, CY, and PM of HEK293 cells treated with or without ACh 10^−4^ M for 5 min. C, Western blotting displaying the IQGAP1, p122RhoGAP/DLC-1, and PLC-δ1 protein level in the TCL, CY, and PM of HEK293 cells with and without IQGAP1 siRNA transfection. D, Double immunostaining of PLC-δ1 and IQGAP1 in skin fibroblasts with ionomycin. The localization of PLC-δ1 is indicated by the red fluorescence of Texas Red-X (a, d, and g). The localization of IQGAP1 is indicated by the green fluorescence of Alexa Fluor 488 (b, e, and h). Merged images are shown (c, f, and i). One flat, extended area is magnified (d, e, f, g, h, and i). Scale bars, 10 μm. E, Double immunostaining of PLC-δ1 and p122RhoGAP/DLC-1 in skin fibroblasts with ionomycin. The localization of PLC-δ1 is indicated by the red fluorescence of Texas Red-X (a and d). The localization of p122RhoGAP/DLC-1 is indicated by the green fluorescence of FITC (b and e). Merged images are shown (c, and f). One flat, extended area is magnified (d, e, and f).

In addition, we assessed whether IQGAP1, p122RhoGAP/DLC-1, and PLC-δ1 were recruited to the PM after stimulation with ACh in HEK293 cells. After the ACh treatment, no change was noted in the amounts of IQGAP1 and p122RhoGAP/DLC-1 in the TCL (Fig 6B) or CY, but both IQGAP1 and p122RhoGAP/DLC-1 displayed clear elevation in the PM. Notably, p122RhoGAP/DLC-1 was not expressed in the cell membrane without the ACh treatment. The PLC-δ1 expression was very low in the TCL and exhibited no change in the CY after the ACh treatment but a marginal increase in the PM. These findings suggested that IQGAP1, p122RhoGAP/DLC-1, and PLC-δ1 moved from the CY to the PM after the ACh treatment in HEK293 cells.

Next, we assessed whether IQGAP1 siRNA was involved in ACh-induced PM recruitment of p122RhoGAP/DLC-1 and PLC-δ1 in HEK293 cells. After transfection with IQGAP1 siRNA, the IQGAP1 protein in the TCL reduced by approximately 60% compared with that in cells transfected with the control vector (Fig 6C); a similar finding was observed in the CY. In addition, marginal amounts of IQGAP1 and p122RhoGAP/DLC-1 were detected in the PM, indicating that a decline in IQGAP1 decreased the recruitment of p122RhoGAP/DLC-1 to the cell membrane in response to ACh.

After 5-min treatment with ionomycin at 10^−5^ M, PLC-δ1 was detected as small dots in the cytoplasm and nucleus (Fig 6D, a). We detected IQGAP1 as small dots in the cytoplasm and nucleus, primarily localized in the nucleus (Fig 6D, b). In addition, small dots of PLC-δ1 and IQGAP1 were colocalized predominantly in the cytoplasm and PMs, where they formed structures analogous to actin stress fibers (Fig 6D, c). For further investigation, we selected two extended flat areas of the cell and combined the PLC-δ1 and IQGAP1 magnified images. These findings suggested that small dots of PLC-δ1 and IQGAP1 were colocalized on a structure that resembled actin stress fibers (Fig 6D, d, e, and f), with a distribution similar to that typical of actin filaments; furthermore, they were colocalized in the PM (arrow; Fig 6D, g, h, and i).

Likewise, we detected PLC-δ1 as small dots in the cytoplasm and nucleus (Fig 6E, a). In addition, p122RhoGAP/DLC-1 was detected as small dots in the cytoplasm and nucleus, but primarily localized in the nucleus (Fig 6E, b). Accordingly, we selected one extended flat area of the cell and combined PLC-δ1 and p122RhoGAP/DLC-1 magnified images. Small dots of PLC-δ1 and p122RhoGAP/DLC-1 were colocalized predominantly in the cytoplasm, again forming structures similar to actin stress fibers; they were also detected in the PMs (Fig 6E, d–f).

### ▪ Regulation of RhoA activation by IQGAP1 and p122RhoGAP/DLC-1

Reportedly, the binding of RhoA–GTP to IQGAP1 regulates RhoA (Casteel et al., 2012), and that p122RhoGAP/DLC-1 exhibits strong GAP activity for RhoA (Homma & Emori, 1995). Thus, we assessed how the overexpression of IQGAP1 and p122RhoGAP/DLC-1 affects RhoA-GAP levels. When HEK293 cells were stimulated with fetal bovine serum (FBS), RhoA-GTP was markedly increased; conversely, the overexpression of p122RhoGAP/DLC-1 and IQGAP1 decreased RhoA–GTP (Fig 7). The amount of GTP bound to the FBS-stimulated Rho protein was elevated by 15.5 ± 3.7 times compared with that of control cells without stimulation (Fig 7B). The amount of GTP bound to the RhoA protein was reduced in p122RhoGAP/DLC-1–transfected cells compared with FBS-stimulated cells in control cells; however, this difference was not statistically significant. When HEK293 cells were transfected with IQGAP1, RhoA activation in FBS-stimulated cells was markedly decreased. In addition, when HEK293 cells were cotransfected with IQGAP1 and p122RhoGAP/DLC-1, RhoA activation was markedly decreased in FBS-stimulated cells. We observed a trend toward lower RhoA–GTP levels in cells cotransfected with IQGAP1 and p122RhoGAP/DLC-1 compared with those transfected with IQGAP1 alone; however, this difference was not statistically significant.

**Fig 7.**
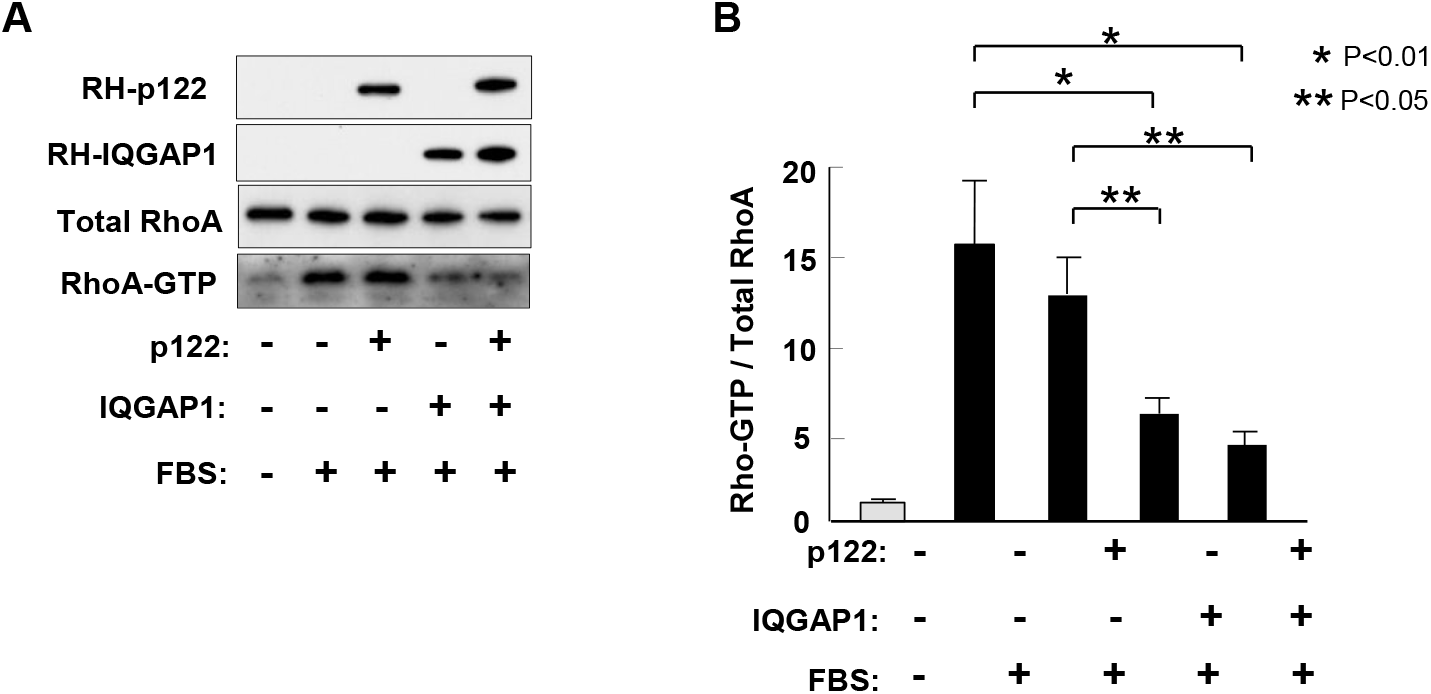
IQGAP1 and p122RhoGAP/DLC-1 regulates the RhoA activation. A, HEK293 cells are subcultured in 6-cm Petri dishes and transfected with pcDNA3/RH-p122, pcDNA3/RH-IQGAP1, or empty vector. Cells are serum-starved for 16 h and stimulated with 20% fetal bovine serum (FBS) for 5 min for the Rho activation. Cell lysates are directly analyzed using Western blotting for p122RhoGAP/DLC-1, IQGAP1, and total RhoA (2% of input). GTP-bound Rho is isolated from cell lysates; RhoA–GTP bound to Rhotekin-RBD is assessed using Western blotting with a RhoA-specific antibody (bottom). B, Western blotting results from three independent experiments performed as in Fig 7A are analyzed by densitometry. The amount of endogenous RhoA–GTP (normalized to total RhoA) present in serum-starved, empty vector transfected cells is assigned a value of 1.

## Discussion

To further our understanding of p122RhoGAP/DLC-1, we searched for partner molecules operating with it in this study. Accordingly, we conducted immunoprecipitation experiments with skin fibroblasts obtained from patients with CSA, using the anti-p122RhoGAP/DLC-1 antibody, and determined that IQGAP1 interacts with p122RhoGAP/DLC-1. In addition, using pull-down experiments, we observed that IQGAP1-C interacts with p122RhoGAP/DLC-1 through the region including the START domain of p122RhoGAP/DLC-1, whereas PLC-δ1 interacts with IQGAP1-N through the region including the CHD and WW domain of IQGAP1-N. Choi et al. (2013) demonstrated by immunostaining that endogenous IQGAP1 was partially colocalized with GFP–PLC-δ1–PH. Yamaga et al. (2008) reported that p122RhoGAP/DLC-1 interacted with the PH domain of PLC-δ1 in a pull-down experiment using various deletion mutants of GST-fused PLC-δ1 as baits. Overall, these findings suggested that p122RhoGAP/DLC-1, IQGAP1, and PLC-δ1 form heteromeric protein complexes.

In this study, we used ACh as a cell stimulant because it is extensively used clinically in provocation tests for coronary spasm (Okumura et al., 1988). The PLC activity and the ACh-induced increment in [Ca^2+^]_i_ were 1.4 times higher in HEK293 cells overexpressing IQGAP1 compared with cells without IQGAP1 transfection. Conversely, the IQGAP1 knockdown led to a diminished [Ca^2+^]_i_ response in HEK293 cells. In addition, cotransfection with IQGAP1 and p122RhoGAP/DLC-1 siRNA to disrupt the complex decreased the [Ca^2+^]i response to ACh in HEK293 cells. These findings suggest that the p122RhoGAP/DLC-1, PLC-δ1, and IQGAP1 complex is vital for the calcium response activation. p122RhoGAP/DLC-1 interacts with the PH domain of PLC-δ1 to enhance its activity and produces, through the hydrolysis of PIP_2_, two second messengers, IP_3_ and diacylglycerol (Homma & Emori, 1995, Yamaga et al., 2008). We found that IQGAP1 interacted with PLC-δ1 and that the IQGAP1 overexpression enhanced the PLC activity. These findings suggest that the direct binding of IQGAP1 to PLC-δ1 is crucial for the activation of PLC-δ1. As p122RhoGAP/DLC-1 binds to the PH domain of PLC-δ1 and activates PLC-δ1, it is essential to ascertain the domain of PLC-δ1 that binds IQGAP1-N, including the CHD and WW domains, in the future study.

IQGAP1 is a ubiquitously expressed multimodular scaffold protein that interacts with various proteins in several cell types (Hedman et al., 2015, Noritake et al., 2005, White et al., 2012). Scaffold proteins facilitate the assembly of signaling cascades by concurrent binding to multiple consecutive components in the signaling pathway; by doing so, they regulate the speed, specificity, intracellular localization, and amplification of signal propagation (Pan et al., 2012). Scaffold proteins for the mitogen-activated protein kinase (MAPK) cascade were among the first to be discovered (Printen et al., 1994, Therrien et al., 1996). The expanding group of MAPK scaffolds comprises several scaffolds for the extracellular signal-regulated kinase (ERK) pathway (Choi et al., 2017), such as kinase suppressor of Ras 1 (KSR1), paxillin, MEK partner 1 (MP1), caveolin-1, and IQGAP1 (Kolch, 2005, Wortzel et al., 2011). Reported target proteins of IQGAP1 are actin, Ca^2+^/calmodulin, E-cadherin, epidermal growth factor receptor, B-Raf, MAPK/ERK kinase1/2, ERK2, and Cdc42 (Pan et al., 2012). Caveolins are a family of integral membrane proteins that constitute the principal components of caveolae membranes and are involved in receptor-independent endocytosis (Scherer et al., 1996, Tang et al., 1996, Williams et al., 2004). The IQGAP1 colocalization with the prescribed caveolae marker protein, caveolin-1, has been established by confocal microscopy and proximity assay (Jufvas et al., 2016). Yamaga et al. (2004) reported that p122RhoGAP/DLC-1 binds caveolin-1 through the RhoGAP domain, and that the START domain could bind to cholesterol. Furthermore, in PC12 cells, the agonist-induced primary increase in Ca^2+^ recruits PLC-δ1 into lipid rafts from other parts of the PM or the CY by other scaffold proteins (Yamaga et al., 2004). In this study, we assessed the ionomycin-induced recruitment of p122RhoGAP/DLC-1, IQGAP1, and PLC-δ1 in skin fibroblasts by Western blotting and fluorescence microscopy, as well as the ACh-induced recruitment of these proteins in HEK293 cells by Western blotting. After treatment with ionomycin and ACh, the amount of p122RhoGAP/DLC-1 and PLC-δ1 in the CY decreased, whereas p122RhoGAP/DLC-1, IQGAP 1, and PLC-δ1 increased in the PM. Nevertheless, how the initial increase in intracellular Ca^2+^ promotes the PM recruitment of PLC-δ1 remains unclear. Contrarily, this study revealed that IQGAP1, p122RhoGAP/DLC-1, and PLC-δ1 are recruited to the PM by ACh and ionomycin stimulation. IQGAP1 is a novel stimulating protein that complexes with p122RhoGAP/DLC-1 and PLC-δ1 to move to the PM and enhance the PLC activity.

Fig 8 shows protein complexes formed by IQGAP1, p122RhoGAP/DLC-1, and PLC-δ1. The PLC-δ1–binding site is located at IQGAP1 between aa residues 1 and 745; notably, this region includes a CHD and WW domain. The IQGAP1-C binds to the C-terminal region of p122RhoGAP/DLC-1 between aa residues 802 and 1091. In addition, this region includes a START domain. As p122RhoGAP/DLC-1 interacts with the PH domain of PLC-δ1, the protein complex shown in Fig 8 was conceived. Yamaga et al. (2008) reported that p122RhoGAP/DLC-1 is constitutively expressed in lipid rafts in rat pheochromocytoma PC12 cells. In addition, PLC-δ1 amount of the lipid raft fraction increased after the carbamylcholine treatment. These findings suggest that p122RhoGAP/DLC-1 and PLC-δ1 might separately form complexes with IQGAP1. Nevertheless, further experiments are warranted. IQGAP1 forms a complex with kinesin Kip2 via Bik1 (CLIP-170), and CLIP-170 captures the plus end of microtubules (Fukata et al., 2002, Noritake et al., 2005). Moreover, fluorescent-labeled CLIP-170 and kinesin Kip2 have been shown to comigrate along individual microtubule (Carvalho et al., 2004). Hence, IQGAP1 in the complex, perhaps, interacts with CLIP-170, which moves along microtubules by interacting with kinesin (Fukata et al., 2002) (Fig. 8).

**Fig 8.**
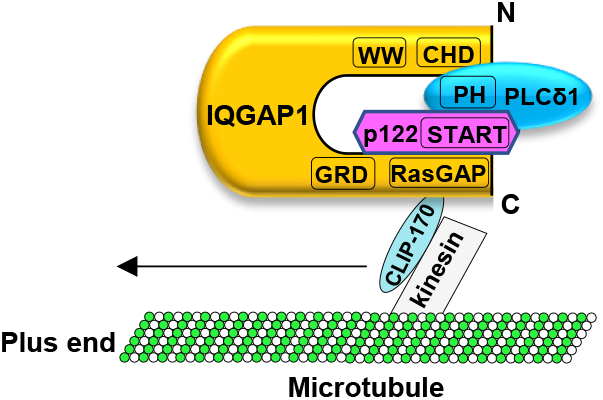
The possible complex formation of IQGAP1, p122RhoGAP/DLC-1, and PLC-δ1.

PLC-δ1 in the lipid rafts “meets” p122RhoGAP to be activated in the presence of Ca^2+^, leading to the robust PIP_2_ hydrolysis and forming clusters in the lipid rafts. Hydrolyzed IP3 and diacylglycerol activate IP3R and transient receptor potential channels and further elevate the influx of Ca^2+^ from voltage-gated calcium channels. Yamaga et al. have investigated the recruitment of p122RhoGAP/DLC-1 and PLC-δ1 to lipid rafts using rat pheochromocytoma PC12 cells. The amount of PLC-δ1 in the lipid raft fractions increased after the carbamylcholine treatment, whereas p122RhoGAP/DLC-1 remained unchanged, suggesting that p122RhoGAP/DLC-1 is constitutively expressed in lipid rafts (Yamaga et al., 2008). Kawai et al. (2004) reported that endogenous p122RhoGAP/DLC-1 was observed at the tips of actin stress fibers. In this study, we investigated the recruitment of IQGAP, p122RhoGAP/DLC-1, and PLC-δ1 to the cell membrane using HEK293 cells and skin fibroblasts. In HEK293 cells, IQGAP1 and p122RhoGAP/DLC-1 clearly increased in the PM after the ACh treatment. Furthermore, in skin fibroblasts, the PM IQGAP1, p122RhoGAP/DLC-1, and PLC-δ1 clearly elevated following ionomycin treatment. We found the amount of p122RhoGAP/DLC-1 in the lipid raft fraction or PM differs based on the type of cells. In this study, immunostaining revealed that PLC-δ1 and IQGAP1 were colocalized on a structure that resembled actin stress fibers, with a distribution analogous to typical actin filaments after the ionomycin treatment in skin fibroblasts. We observed that the initial increase in intercellular Ca^2+^ promoted the recruitment of IQGAP1, p122RhoGAP/DLC-1, and PLC-δ1 to the cell membrane. Nevertheless, how these molecules are released from microtubules and recruited to PMs or lipid rafts remains unclear, necessitating the elucidation of underlying mechanisms. The protein complex of IQGAP1, p122RhoGAP/DLC-1, and PLC-δ1 increased in cells obtained from patients with CSA compared with those from control subjects, being associated with the enhanced PLC activity. Hence, IQGAP1 could be a novel stimulator for the PLC-δ1 activity through direct activation and transporting to the PM and may, thus, play a pivotal role in the CSA pathogenesis.

The p122RhoGAP/DLC-1 protein performs another function, as a GAP activator for Rho. Reportedly, the Rho-kinase inhibitor, fasudil, attenuated the constrictor response of the coronary artery to ACh and prevented the occurrence of chest pain in patients with CSA (Masumoto et al., 2002). Through its GAP activity, p122RhoGAP/DLC-1 might antagonize the development of coronary spasm similar to the Rho-kinase inhibitor. Casteel et al. (2012) reported that IQGAP1 also functions against Rho. IQGAP1-C (aa 863–1657) binds directly to RhoA; when 293T cells were cotransfected with epitope-tagged RhoA and IQGAP1, the amount of GTP bound to the Rho protein almost doubled compared with control cells transfected with RhoA plus an empty vector. When MDA-MB-231 cells were transfected with an siRNA specific for IQGAP1, the IQGAP1 depletion markely declined the RhoA activation in FBS-stimulated cells (Casteel et al., 2012). Conversely, Bhattacharya et al. (2014) reported that RhoA–GTP, the active form, was increased in human airway smooth muscle cells with the shRNA-mediated IQGAP1 knockdown. Some prior studies reported an elevation in the p122RhoGAP/DLC-1 protein expression in cultured skin fibroblasts obtained from patients with CSA (Murakami et al., 2010). In addition, the IQGAP1 protein expression was increased in cultured skin fibroblasts obtained from patients with CSA in this study. We assessed the RhoA activity and the elevation in intracellular free Ca^2+^ using IQGAP1-transfected cells. Transfection with p122RhoGAP/DLC-1 decreased the RhoA activity compared with FBS-stimulated controls, and transfection with IQGAP1 further strongly decreased the RhoA activity. In addition, cotransfection with p122RhoGAP/DLC-1 and IQGAP1 considerably decreased the RhoA activity compared with FBS-stimulated controls, and its activity approached the level of unstimulated control HEK293 cells. These findings suggest that the p122RhoGAP/DLC-1 and IQGAP1 overexpression exerted a synergistic effect on the RhoA reduction. Overall, these findings suggest that the Rho-GAP cascade might compensate enhanced coronary spasm but not contribute to enhanced coronary spasm.

## Materials and Methods

### Patients

This study protocol was approved by the Ethics Committee of our institution, and we obtained written informed consent from all patients before the study. Our study cohort comprised 8 Japanese patients with CSA (7 male and 1 female; mean age: 58 ± 6 years) and 6 control subjects without hypertension or any history suggestive of angina pectoris (3 male and 3 female; mean age: 52 ± 6 years). All patients with CSA underwent coronary arteriography with an intracoronary administration of ACh to induce coronary spasm, defined as the total or subtotal occlusion or severe vasoconstriction of the coronary artery related to chest pain and ischemic change on ECG. After an intracoronary injection of isosorbide dinitrate, coronary arteriograms revealed normal or almost normal coronary arteries with diameter stenosis ≤50% of the lumen diameter in all patients with CSA.

### Cell culture

We prepared human skin fibroblasts using the explant method, as described previously (Okumura et al., 2000). The fibroblasts and HEK293 cells were cultured in Dulbecco’s modified Eagle’s medium (#11965–092; Gibco, NY) supplemented with 10% FBS (#12483-020; Gibco), penicillin (100 U/mL), and streptomycin (100 μg/mL; #15140-122; Gibco) at 37°C under 5% CO_2_ and 95% air.

### Immunoprecipitation

We extracted skin fibroblasts with Pierce IP Lysis Buffer (87787; ThermoFisher, MA) containing a protease inhibitor cocktail (78429; ThermoFisher). The cell debris was removed from the lysates by centrifuging at 13,000 rpm for 15 min at 4°C, followed by preclearing with Dynabeads G (DB10003; Dynal Biotech, Oslo, Norway). We added the Dynabeads G (50 μL) to the anti-p122RhoGAP/DLC-1 antibody (612020; BD Biosciences, NJ), anti-IQGAP1 antibody (610611; BD Biosciences), or normal mouse IgG, incubated for 2 h at 4°C, and washed with phosphate-buffered saline (PBS) with 0.02% Tween 20. Next, the cell lysates were added and incubated overnight at 4°C.

After incubation, we washed the immunoprecipitates three times with PBS with 0.02% Tween 20 and eluted the immobilized immunocomplexes with a sample-treating solution containing 2% SDS and 5% β-mercaptoethanol for 30 min at 50°C. Then, the eluted proteins were fractionated by SDS–PAGE and detected with a Silver Stain MS Kit (299–58901; FUJIFILM Wako Pure Chemical Corporation, Osaka, Japan) or by Western blotting.

### Mass spectrometry analysis

We analyzed protein bands (molecular weight range: 150–250 kDa) that interacted with the p122RhoGAP/DLC-1 peptide in an immunoprecipitation assay using skin fibroblasts by TOF-MS (Oncomics Co., Ltd.).

### Plasmid construction and transfection

The cDNA for human pEGFP (enhanced green fluorescent protein)-C2-IQGAP1, pGEX-IQGAP1-N (1–863), and pGEX–IQGAP1-C (746–1657) were kindly provided by Dr. Kozo Kaibuchi (Department of Cell Pharmacology, Nagoya University Graduate School of Medicine, Nagoya, Japan). To express p122RhoGAP/DLC-1, the cDNA was amplified by a polymerase chain reaction from a human brain cDNA library (Clontech, CA) using an appropriate primer pair. To express proteins tagged with an epitope at the N terminus in human cells, we inserted the IQGAP1 and p122RhoGAP/DLC-1 cDNA into the plasmid vector pcDNA3/RH-N (Kamitani et al., 1998) to create pcDNA3/RH-IQGAP1 and pcDNA3/RH-p122RhoGAP/DLC-1. We subcloned cDNA fragments corresponding to each p122RhoGAP/DLC-1 fragment into pTrcHisA to obtain p122RhoGAP/DLC-1 (aa 1–1079), p122RhoGAP/DLC-1 (aa 1–546), p122RhoGAP/DLC-1 (aa 1–801), and p122RhoGAP/DLC-1 (aa 547–1079). Furthermore, pEGFP-C2-IQGAP1 and pcDNA3/RH-p122RhoGAP/DLC-1 plasmids were transfected into HEK293 cells using Lipofectamine 3000 (Invitrogen, MA).

### RNA interference

HEK293 cells were transfected with p122RhoGAP/DLC-1 siRNA (sense, 5′-GAAACGCCUUAAGACACUATT-3′; antisense, 5′-UAGUGUCUUAAGGC GUUUCTT-3′; TaKaRa Biotechnology Co., Ltd., Kyoto, Japan), IQGAP1 siRNA (s16837; Applied Biosystems), or negative control. The cells were transfected by siRNA (final concentration: 100 nM) at 70%–80% confluency using a transfection reagent, DharmaFECT Duo (T-2010-03; Dharmacon, CO), in the complete medium, per the manufacturer’s instructions.

### Western blotting

We treated the whole-cell protein samples for 30 min at 50°C in a sample-treating solution containing 2% SDS and 5% β-mercaptoethanol. We followed the protocol provided for the Plasma Membrane Protein Extraction Kit (BioVision, CA) to extract the PM protein. Next, protein samples were separated by SDS-PAGE and electrophoretically transferred to a polyvinylidene fluoride membrane (Bio-Rad Laboratories, Hercules, CA). After 1-h blocking, we incubated the membranes overnight at 4°C with the primary antibodies for IQGAP1, p122RhoGAP/DLC-1, PLC-δ1 (ab134936; Abcam, Cambridge, UK), and GAPDH (sc-25778S; Santa Cruz Biotechnology, TX). We used a horseradish peroxidase–conjugated antibody (Santa Cruz Biotechnology) as a secondary antibody. After SDS–PAGE, we performed Western blotting per the protocol provided with the ECL (Enhanced Chemiluminescence) Detection System (GE Healthcare, IL). Furthermore, we performed densitometric analysis with Scion imaging software and evaluated the relative ratio to GAPDH for each sample.

### Fluorescence microscopy

We performed fluorescence microscopy studies to investigate the subcellular locations in skin fibroblasts of IQGAP1, p122RhoGAP/DLC-1, and PLC-δ1. Skin fibroblasts were cultured on a 3.5-cm glass-bottom dish; the cells were fixed in a 4% paraformaldehyde solution (pH 7.5) for 15 min and permeabilized with 0.1% Triton X-100 for 15 min at room temperature. Next, the cells were labeled with one of the following primary antibodies overnight at 4°C: goat polyclonal anti-IQGAP1 (sc-8737; Santa Cruz Biotechnology), rabbit polyclonal anti-p122RhoGAP/DLC-1 (sc-32931; Santa Cruz Biotechnology), mouse monoclonal ant-PLC-δ1 (sc-374329; Santa Cruz Biotechnology), or rat monoclonal anti-tubulin (ab6160; Abcam). After washing, the cells were labeled with Alexa Fluor 488 rabbit anti-goat IgG (H+L), Texas Red-X–conjugated goat anti-rabbit IgG, Alexa Fluor 594–conjugated rabbit anti-goat IgG (H+L), FITC-conjugated goat anti-mouse IgG, or Alexa Fluor 488-conjugated donkey anti-rat IgG (H+L) antibody (ThermoFisher) at a dilution of 1:1000 for 1 h at room temperature. Finally, we analyzed the cells with a BZ-X700 fluorescence microscope (Keyence, Osaka, Japan).

### Measurement of [Ca^2+^]_i_

We subcultured HEK293 cells in 6-cm Petri dishes and transfected with human IQGAP1 cDNA or an empty vector (3.0 μg DNA/well for all). After loading with 5 μmol/L Fura-2 AM, ACh at 10^−4^ M was added, and we measured the [Ca^2+^]_i_ response at excitation wavelengths of 340 and 380 nm and an emission wavelength of 510 nm, as described previously (Osanai et al., 1992). We used ACh because it is extensively used to induce coronary spasm in Japanese patients (Okumura et al., 1988). Furthermore, calibration was performed using ionomycin followed by EGTA–Tris.

### Measurement of the PLC activity

The PLC assay system comprised the following components: N-2-hydroxyethylpiperazin-N’-2-ethanesulfonic acid (50 mmol/L); calcium chloride (0.1 mmol/L); sodium cholate (9 mmol/L); tritium-PIP_2_ (40,000 counts/min); and the cell protein (100 μg). The reaction was discontinued with a combination of chloroform, methanol, and hydrogen chloride, followed by 1-N hydrogen chloride containing EGTA. After extraction, we removed the aqueous phase for liquid scintillation counting.

### *In vitro* interaction assay

Several recombinant proteins were first expressed in *Escherichia coli* (BL21 DE333) using the eukaryotic expression vectors pGEX- and pTrcHisA-, as described previously (Kaelin et al., 1991, Kamitani et al., 1997). Then, GST-fused IQGAP1-N and IQGAP1-C were purified using Glutathione Sepharose 4B (17-0756-01; GE Healthcare). Next, we centrifuged the bacterial crude lysates containing p122RhoGAP/DLC-1 (aa 1–1079), p122RhoGAP/DLC-1 (aa 1–546), p122RhoGAP/DLC-1 (aa 1–801), p122RhoGAP/DLC-1 (aa 547–1079), and PLC-δ1 (1–756) at 14,000 × *g* for 5 min and incubated the supernatants for 3 h at room temperature with GST fusion proteins immobilized on Glutathione Sepharose beads. Then, the beads were washed six times with the lysis buffer, and the proteins precipitated on the beads were eluted in 2% SDS treating solution. Finally, we subjected them to Western blotting using a mouse anti-RH monoclonal antibody (specific to the aa sequence RGSHHHHHH).

### Localization of PLC-δ1, IQGAP1, and p122RhoGAP/DLC-1

We cultured HEK293 cells in 10-cm culture plates at a density of 5 × 10^6^ cells/dish. After 24-h incubation, the cells were serum-starved for 16 h and stimulated with ACh (10^−4^ M) for 5 min. We cultured skin fibroblasts and stimulated them with ionomycin (10^−5^ M) for 5 min. Then, we extracted CY and PM proteins using the PM Protein Extraction Kit (BioVision). The protein samples were treated for 30 min at 50°C in a sample-treating solution containing 2% SDS and 5% β-mercaptoethanol, followed by subjecting them to Western blotting. Furthermore, we assessed the localization of PLC-δ1, IQGAP1, and p122RhoGAP/DLC-1 by a cell preparation method similar to fluorescence microscopy.

### RhoA-GTP pull-down assay

We measured the RhoA activity using an Active Rho Detection Kit (Cell Signaling Technology, MA), per the manufacturer’s protocol. Then, HEK293 cells were subcultured in 6-cm Petri dishes and transfected with pEGFP-C2-IQGAP1, pcDNA3/RH-p122, or an empty vector (1.0 μg DNA/well for all). Finally, the cells were serum-starved for 16 h and, then, stimulated with 10% FBS for 5 min to activate RhoA.

### Statistical analysis

In this study, data were analyzed using the statistical software JMP (version 11.0) and were expressed as mean ± standard deviation. We tested comparisons of two variables using paired or unpaired *t*-tests, as appropriate, as well as multiple comparisons using the Tukey–Kramer test. Of note, *P* < 0.05 was considered statistically significant.

## Acknowledgments

We thank Dr. Kozo Kaibuchi for kindly providing human pEGFP-C2-IQGAP1, pGEX-IQGAP1-N cDNA, and pGEX-IQGAP1-C cDNA. Drs. Hirofumi Tomita and Kenji Hanada have received research grant support from JSPS KAKENHI (grant numbers: JP16K09489 and JP18K15875), and Hirofumi Tomita has received research grant support from The Uehara Memorial Foundation.

